# Electrocorticographic Network Feature Space Constriction as a Preictal Biomarker

**DOI:** 10.64898/2026.07.08.736809

**Authors:** Jeremy Goetz, John Beggs, Robert Worth, Louis R. Nemzer

**Affiliations:** Department of Physics, Indiana University, Bloomington, USA; Department of Mathematical Sciences, Indiana University School of Medicine, Indianapolis, USA; Halmos College of Arts and Sciences, Nova Southeastern University, Fort Lauderdale, USA

## Abstract

In patients with epilepsy, seizures are associated with pathological neural synchronization. However, the preictal period preceding a seizure often exhibits reduced spatial synchronization compared to normal cognition. This observation aligns with the concept of the brain as a complex dynamical system, where a reduction in dimensionality and resilience can precede a phase transition. The Critical Brain Hypothesis suggests a connection between the loss of healthy scale-free behavior and various disorders, including epilepsy. Our study investigates preictal changes by utilizing network features, such as mean node degree and mean clustering coefficient, derived from thresholded correlation matrices of patient intracranial electrocorticographic electrode data. We observed a suppression of intermittent high-synchronization periods within the feature space during the minutes leading up to seizure onset. This constriction of the explored hypervolume in the preictal state indicates a breakdown in the brain’s ability to maintain normal coherence. We use these preictal changes to predict the probability of seizure onset using a Support Vector Machine algorithm. These discrete predictions can then be combined into real-time continuous seizure risk forecasts via Bayesian updating. This innovative and computationally lightweight approach has the potential to significantly improve upon static predictions, providing opportunities for more adaptable, quantitative, and interpretable tools for managing seizures.

## INTRODUCTION

### Motivation

Epilepsy impacts approximately 1% of people worldwide, making it one of the most common chronic neurological conditions. Characterized by recurrent seizures caused by pathological electrical brain activity, about 30% of epilepsy patients do not find their symptoms controlled by medication (Chen et al., 2018). For patients with medication-refractory focal epilepsy, resective surgery does not achieve freedom from disabling seizures in approximately 30-40% of cases (Englot & Chang, 2014). The unpredictable nature of seizure onset can prevent these individuals from fully participating in activities, leading to a significantly reduced quality of life (L. R. Nemzer et al., 2020). Some people are precluded from participating in normal activities, such as driving or operating machinery, and the ability to forecast the real-time risk of seizures (Mormann et al., 2005) (Mormann et al., 2007) for patients with epilepsy could represent a marked improvement in daily satisfaction, the ability to function independently, and a decrease in hospital admissions (Nass et al., 2016). If it were possible to predict the onset of a seizure even a short time before it happens, it would improve the ability of patients and caregivers to manage the condition and allow for timely therapeutic interventions (Gadhoumi et al., 2016) (Freestone et al., 2017), which are crucial for averting seizures or mitigating their effects. Theoretically, interventions like electrical stimulation (Foutz & Wong, 2022) or prompt administration of medication could prevent the onset of a seizure. A wearable device that continuously monitors biological signals might be able provide patients with warnings during times of elevated risk (Nasseri et al., 2021).

### Critical Brain Hypothesis

Healthy brain function is believed to operate at or near a critical state maintained as a homeostatic setpoint (Hengen & Shew, 2025), and characterized by power-law behavior (Beggs & Plenz, 2003) (L. Nemzer, 2019). This level of activity, balanced between chaos and stasis, optimizes information transmission between distant regions and sensitivity to external stimuli (Beggs, 2008), and reflects the brain’s ability to process data and adapt to variable conditions. We suggest that the preictal state just prior to a seizure reflects a deviation from this balance, characterized by a constriction of phase space hypervolume explored and suppression of rare neural events. Recent experiments have linked absence seizures to the confinement of dynamics on an attractor that relies on the same circuit substrate associated with the loss of consciousness when using general anesthesia (Hull et al., 2025). The scale-free nature (He, 2011) of power laws dictate that the brain normally has the capability to explore even distant regions of feature space, with excursions representing the long tail of the distribution. Healthy cognition therefore consists of a wide range of activity, traversing a large hypervolume of the available phase space. The trajectory in phase space would therefore be complex and unpredictable, reflecting a high-dimensional system.

### Paradoxical Desynchronization

Seizures represent a state of pathological synchronization, in which nodes of a network can be recruited to combine in aberrant activity (Boddeti et al., 2025) like the nucleation of a crystal. Somewhat paradoxically, the time period leading up to a seizure may exhibit desynchronization (Mormann et al., 2003) (Netoff & Schiff, 2002) and a simplification of brain dynamics. This process can be accompanied by a decrease in the dimensionality of the EEG signal and a reduction in the hypervolume of the phase space explored. As a result, the preictal state loses resilience to perturbations (Chang et al., 2024). This desynchronization has been well-documented in human intracranial and extracranial recordings, and was observed in both neural activity and local field potentials, *in vitro* and *in vivo* (Jiruska et al., 2013). Leading up to a seizure, the low-complexity state creates a vulnerable condition in which the system is more easily drawn into the attractor state that defines the seizure (E. T. Wang et al., 2022). These findings suggest a two-stage process. First, an initial functional decoupling of distant brain regions occurs at the onset of a seizure, followed by an unusually strong re-coupling as the seizure progresses (Wendling et al., 2003). Other pathological states may be indicated by a simplification of physiological signals. For instance, diminished heart rate variability can suggest subclinical cardiac disease. (Tsuji et al., 1996).

### Preictal Period

Despite being central to the question of seizure prediction, the preictal period – which is the hypothesized period of time during which neurological or physiological changes heralding a seizure could be detected in principle – remains poorly understood (Kuhlmann et al., 2018). Some practitioners use an arbitrary threshold, such as 15 minutes prior to seizure onset, as a working definition. However, this empirical approach to labeling seizure events may lead to tautological definitions. If seizures represent critical phase transitions of a dynamical system (L. R. Nemzer et al., 2020), then it is possible that previously recognized phenomena, like critical slowing down, can be identified (Liu et al., 2023; Maturana et al., 2020). Quantifying the preictal period in a rigorous way (Bandarabadi et al., 2015) would not only allow for an assessment of the intrinsic “predictability” of seizures (Mormann et al., 2005), but may also shed light on the underlying mechanism of ictogenesis. The horizon for prediction can be limited both by the sensitivity of monitoring devices, as well as the inherently chaotic nature of nonlinear dynamical behavior. For example, the exponential growth of uncertainty over time is thought to be governed by the largest Lyapunov exponent (Vannitsem, 2017), which for weather forecasting has been measured to correspond to a doubling time of approximately three days (Krishnamurthy, 2019). In the case of seizure prediction, the separation of nearby trajectories would tend towards a practical upper bound on the predictability. The fundamental premise of seizure prediction hinges on the existence of a preictal period, though its level of consistency across patients and seizure types remains mostly unknown. For this study, we conceptualize the preictal period as the interval preceding a seizure when discernible correlations in brain activity can signal an elevated probability of an impending seizure. We build on the idea that seizure onset reflects a network dysfunction (Frassineti et al., 2024), which is evident as a phase transition in electrical brain signals.

## BACKGROUND

### Previous Efforts

Electrical brain monitoring, starting with electroencephalography (EEG), is now a century old (Tudor et al., 2005) (Ojha, 2024). The fusion of modern machine-learning techniques with sophisticated intracranial electrocorticographic (ECoG) brain recording technology has recently generated optimism about overcoming the unpredictable nature of seizures. Efforts toward seizure prediction have involved the analysis of features derived from the time domain, frequency domain, entropy, wavelets, and network-level characteristics (Z. Wang et al., 2024). However, despite many years of effort, no clinical devices for assessing real-time risk have yet achieved widespread adoption. This is largely due to insufficient accuracy when making predictions using unseen data. Various machine-learning approaches have been applied (Zurdo-Tabernero et al., 2023). A recent review of deep-learning seizure prediction articles (X. Zhang et al., 2024) found that training data may not reflect real-world conditions, and as a result, fail to generalize for practical applications. Models trained on one dataset may not work well on others (Ren et al., 2022). Another review (Cherian & Kanaga, 2022) provided a comprehensive overview of the latest research on seizure prediction. Often, high claimed classification accuracies, sometimes exceeding 90% even as much as an hour prior to a seizure, may be illusory due to data leaking from slowly varying noise (West et al., 2023), raising doubts about the clinical usability of many reported prior results. Moreover, the non-stationarity of seizure dynamics and variable preictal duration continue to hinder clinical application (Kalousios et al., 2025). Seizure prediction models are typically developed using patient-specific data, leading to poor transferability across different individuals, since the mechanisms that lead to seizures may vary between patients (Freestone et al., 2017). Even with many seizure prediction models considered state-of-the-art, improvement over chance (IoC) was achieved for less than a third of the patients. (Costa et al., 2024). Models that claimed high performance metrics frequently failed when subjected to rigorous evaluation.

This was often due to inflated or misleading claims, data leakage between training and testing sets, optimizing parameters based on test data, creating artificial correlations, inconsistent metric definitions, not including standard comparison metrics, failing to compare against chance, or using biased sampling of data segments that were too easily distinguished. More generally, older linear models sometimes outperform newer, more complex deep learning models in classification tasks (Lütjens et al., 2025). For comparison, using data from 37 patients in the EPILEPSIAE database, the reported sensitivity was up to 0.75, with about one false positive per hour (Pontes et al., 2024). Using the publicly available CHB-MIT dataset (SWE LOO), the claimed sensitivity was 0.7534, with 4.79 false positives per hour (Ali et al., 2024). Other methods used convolutional neural networks (CNN) on a matrix calculated from the pairwise Magnitude Squared Coherence of electrode channels at various frequencies (Lu et al., 2025) or the synchronization of theta waves (Schubert et al., 2025).

### Network Features of Functional Connectivity

In addition to the physiological connections between neurons, the brain can be understood based on the functional connectivity between distant regions computed using temporal correlations in activity. The functional connectivity network, along with associated graph theoretical metrics such as node degree distribution and mean clustering coefficient (Bullmore & Sporns, 2009), has been shown to change during the preictal state (Q. Zhang et al., 2020). These changes can be quantified using network features (Akbarian & Erfanian, 2020) on a thresholded graph, in which network nodes are considered connected by an edge if their temporal correlation exceeds a certain threshold value (Q. Zhang et al., 2020). Somewhat like Topological Data Analysis (TDA), which is the study of how geometric entities are “connected” without regard to their absolute orientation (Upadhyaya et al., 2023), network features can robustly capture essential properties of very high dimensional data not easily accessible through conventional methods. Our study focuses on network-level features to understand seizures as failures of network dynamics. In contrast with alternative methods, such as convolutional neural networks (Priya Prathaban & Balasubramanian, 2021) trained on brain recordings converted into images via spectrogram or wavelet transform approaches, network features require significantly less computing power in terms of memory and storage space. These measures are selected because they effectively quantify the network’s tendency toward localized, simple connectivity (clustering coefficient) and overall density (node degree), serving as robust proxies for complexity and dimensionality.

## METHOD

### Network Feature Workflow

We employed expert-annotated intracranial ECoG data obtained from the EPILEPSIAE European Epilepsy Database (Ihle et al., 2012). This resource contains continuous surface and invasive recordings from a total of 275 patients, including 58 with intracranial recordings for up to 125 channels at up to 2,500 Hz sampling frequency (Klatt et al., 2012). The total recording length in this database exceeds 40,000 hours, with an average recording length across the entire cohort of 165 hours, and as many as 500 hours for some patients. This database also contains extensive metadata, including the 3D-coordinates of implanted electrode positions, along with patient information such as age, sex, seizure etiology, and sampling frequency. Seizure events are expertly annotated with both temporal and spatial specifications, along with seizure type. We constructed network features from the correlation matrix of intracranial electrocorticographic (ECoG) channels in patient data, which were then converted into distance metrics. We used the pairwise Pearson correlation matrix ρ_ij_ (Kuang et al., 2024) between preselected electrodes transformed into a natural distance metric 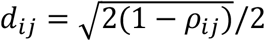 to process the data via established network analysis methods. While correlation ranges from −1 to +1, distance ranges from 0 to 1, with zero representing perfect correlation. Having access to ECoG intracranial patent recordings (Hill et al., 2012) with high temporal and spatial resolution provides significant advantages over publicly available data (Shoeb, 2010), which often consists of scalp EEG recordings that can suffer from artifacts and limited frequency range. In addition, the larger number of patients, as well as the increased number of electrode channels per patient, permits the use of mathematical techniques that would otherwise not be tractable. The workflow is shown in Figure 1.

**Figure 1.**
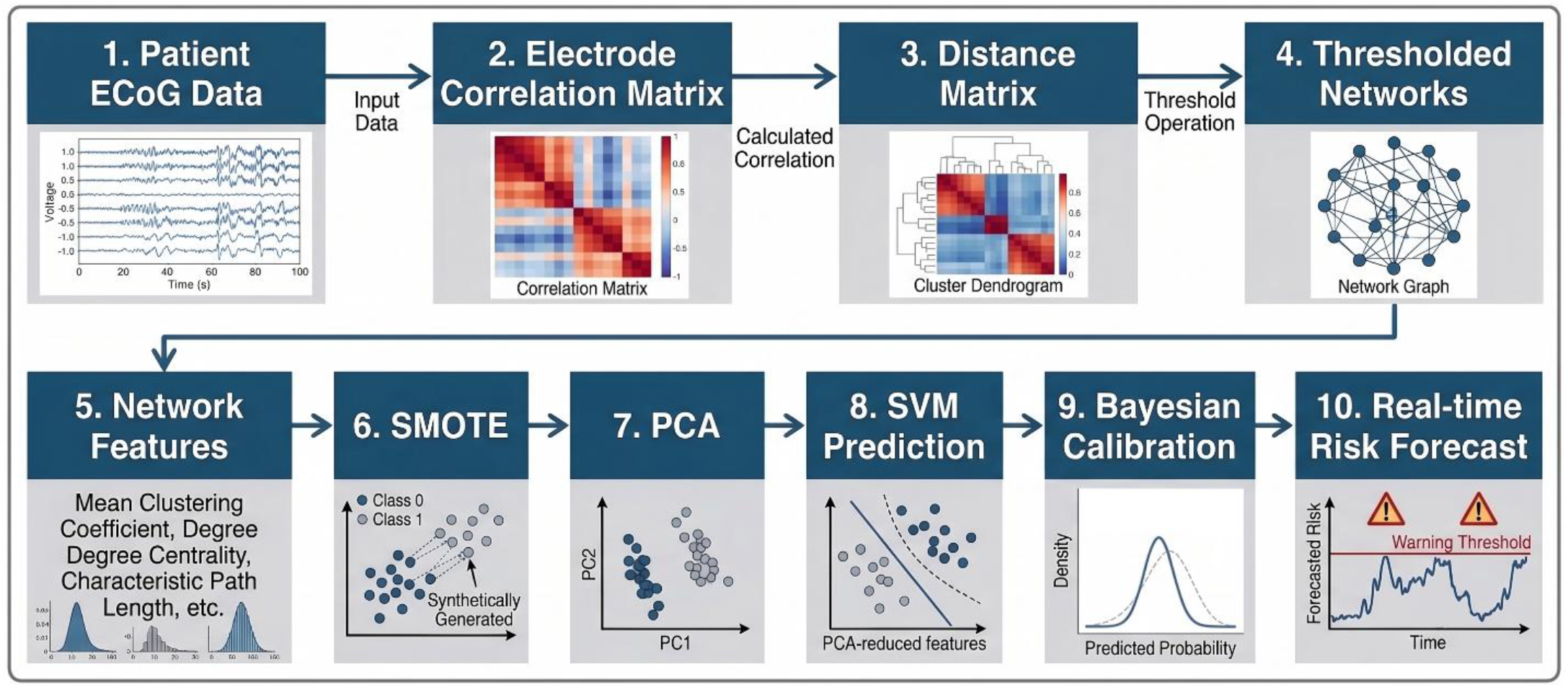
Schematic representation of the analysis workflow. ECoG = electrocorticographic, SMOTE = Synthetic Minority Over-Sampling Technique, PCA = Principal Component Analysis, SVM = Support Vector Machine.

Our seizure prediction algorithm, designed for high-precision temporal analysis, was configured to use 64 electrode channels. Data is stored in one-hour blocks, and recordings were filtered to include only those sampled at 1,024 Hz, ensuring a consistent and high-resolution basis for subsequent computations. Expert annotation indicated the onset time of seizures, and for this analysis, only blocks that contained a seizure were selected for analysis. The data points were classified into “core preictal,” (1-4 minutes prior to seizure onset), intermediate preictal (4-10 minutes) and interictal (10-59 minutes). As discussed below, empirical testing showed that prediction accuracy was improved when intermediate preictal data was masked and not included in training. The cutoff of four minutes was also empirically determined to optimize predictive accuracy. Data less than one minute before seizure onset was completely excluded. Machine learning testing was done on held-out seizure events not included in the training dataset. The analysis used dynamic functional connectivity (Joyce et al., 2010), assessed through the construction of a time-varying correlation matrix. This matrix was computed from the synchronized voltage traces over rolling five-second windows, advancing with a one-second stride. This overlap allows for smooth, continuous tracking of network topology evolution. Example data reflecting an interictal (Figure 2) and preictal (Figure 3) case are shown. Each node represents a different ECoG channel, and edges between nodes become increasingly filled in as the distance threshold is raised.

**Figure 2.**
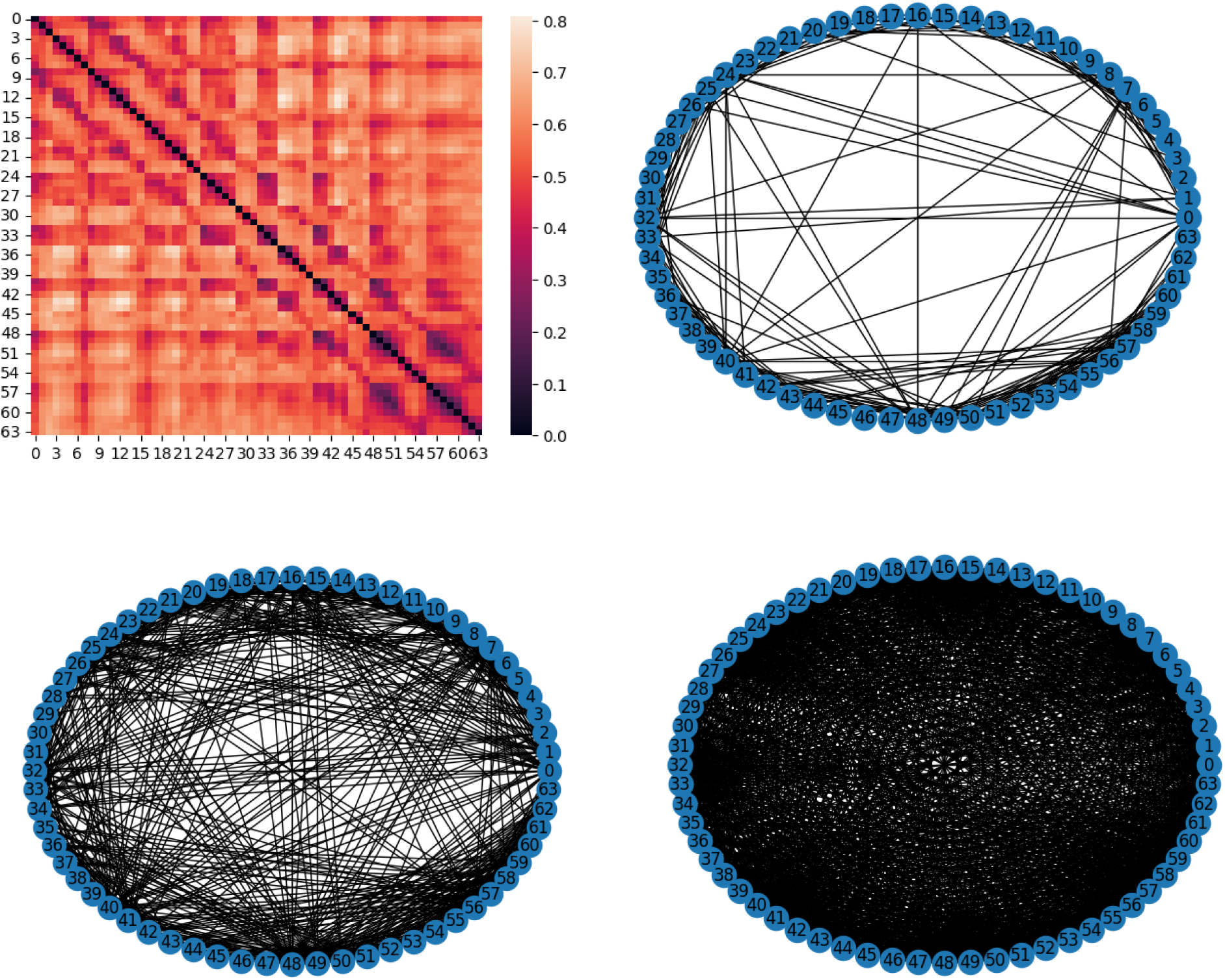
(Clockwise from upper left) Example interictal distance matrix and corresponding thresholded networks at [0.4,0.5, or 0.6] distance thresholds for an interictal data point.

**Figure 3.**
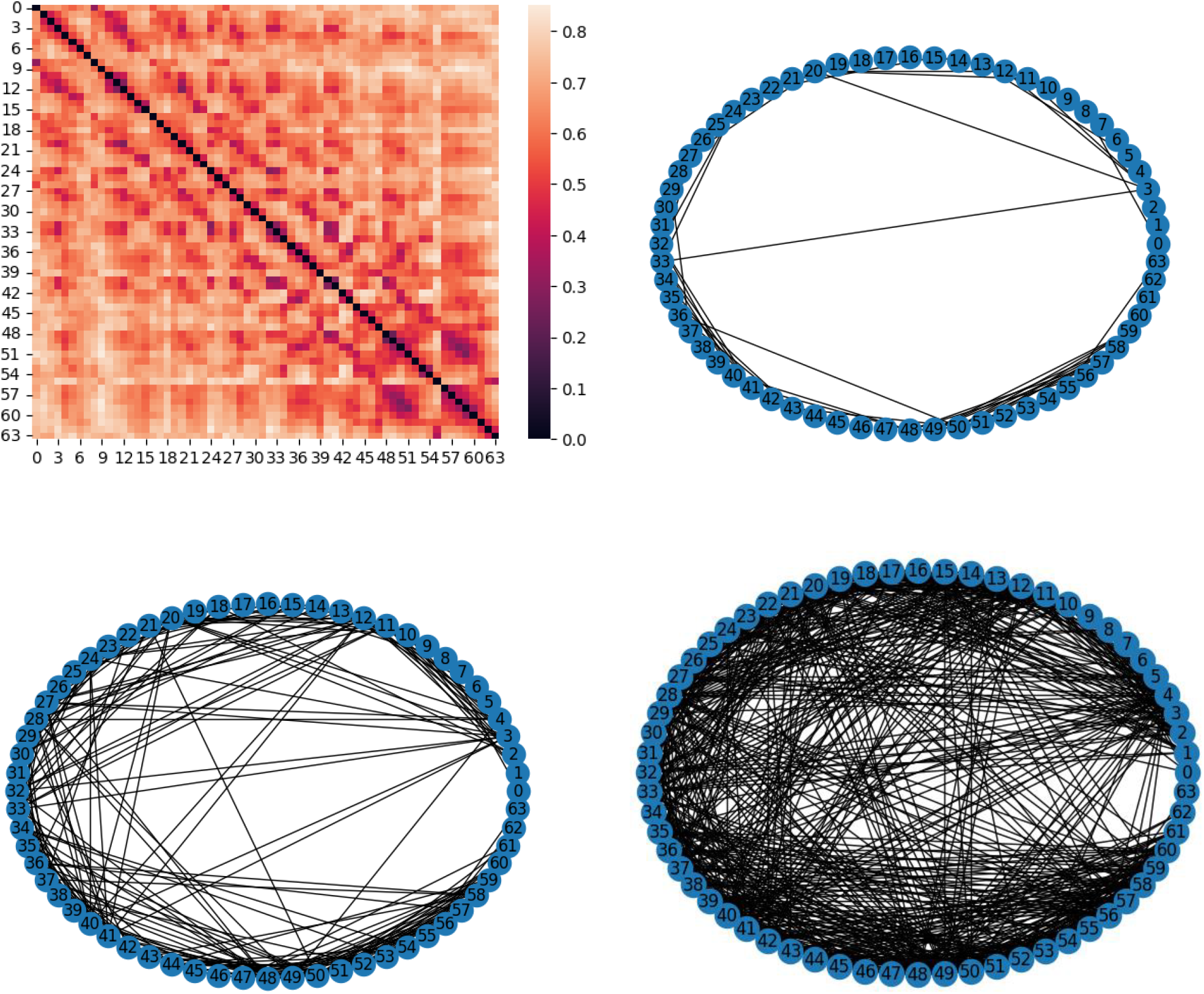
(Clockwise from upper left) Example interictal distance matrix and corresponding thresholded networks at [0.4,0.5, or 0.6] distance thresholds for a preictal data point.

Four established network features were computed for analysis at each threshold, derived from the distance matrix which serves as a representation of the functional brain network. Here, node degree is defined as the number of edges connected to a given node:

1. **Normalized Node Degree Variance:** Calculated as the ratio of the Node Degree Variance to the Node Degree Mean, this metric captures the heterogeneity and centralization of the network connectivity.
2. **Size of Largest Clique:** This feature quantifies the maximum size of a fully connected subgraph, reflecting the robustness and integration of highly synchronous local clusters.
3. **First Eigenvalue:** The magnitude of the largest eigenvalue serves as a global measure of network interconnectedness and synchronization.
4. **Average Clustering Coefficient:** This metric measures the local cliquishness of the network, providing an estimate of the density of connections within the neighborhoods of individual nodes.

Eleven distinct correlation threshold levels were systematically applied. This array of thresholds, spaced at 0.025 increments, was: **[0.250, 0.275, 0.300, 0.325, 0.350, 0.375, 0.400, 0.425, 0.450, 0.475, 0.500]**. By calculating each of the four network features at every one of these eleven threshold values, a comprehensive feature vector consisting of 4 x 11 = 44 unique features was constructed for every one-second stride. This multiple threshold approach is important, as it provides the machine learning algorithm with a robust, multiscale representation of the network’s topological characteristics, enabling the detection of subtle shifts in functional organization that may precede seizure onset.

### Machine Learning Data Preprocessing

Forecasting the risk of rare events, such as epileptic seizures, using machine learning models presents a challenge due to inherent data imbalances. In this context, the minority class, defined as the *preictal* state (the period immediately preceding a seizure), is vastly outnumbered by the majority class, the *interictal* state (the period between seizures). This severe class imbalance can lead to models that are heavily biased toward the majority class, resulting in poor predictive performance, particularly in terms of sensitivity. To effectively address this critical data challenge, a Synthetic Minority Over-Sampling Technique (SMOTE) was implemented. SMOTE operates by creating synthetic samples for the minority class, rather than simply duplicating existing examples. By interpolating between neighboring minority class instances in the feature space, SMOTE was used to achieve a balanced representation of both the preictal and interictal classes within the training dataset. Following the dataset balancing, Principal Component Analysis (PCA) was employed to manage the high dimensionality of the feature space. PCA is a linear dimensionality reduction technique that transforms the data into a new set of orthogonal variables, known as principal components (PCs), which retain most of the variance present in the original data. Through systematic analysis, six principal components (PCs) were ultimately retained. This selection aimed to strike an optimal balance, significantly reducing the feature space complexity while preserving sufficient informational content needed for the subsequent seizure prediction.

### Support Vector Machine

We employed a Support Vector Machine (SVM) algorithm to generate probabilistic predictions for individual five-second data points of ECoG data. While the basic operation of an SVM is based on binary classification, distinguishing strictly between two classes, the Support Vector Classifier (SVC) implementation in this project utilized the distance from the decision boundary to extract more information. Specifically, the model was calibrated using Platt scaling to produce an output that represents a confidence score, ranging continuously from 0 to 1. This score directly corresponds to the probability that the specific data point belongs to the preictal class. A value close to 1 indicates a very high confidence in an impending seizure, while a value near 0 suggests the data point is strongly classified as interictal or normal baseline activity. This probabilistic output is essential for transitioning from a simple binary prediction to a calibrated risk-assessment framework.

### Bayesian Updating

The ultimate goal is to transform the stream of discrete *predictions* from incoming individual data points into a real-time updated *forecast* of seizure risk. This forecast should rigorously and quantitatively (Houle et al., 2021) integrate all observations made up to the present moment, effectively bridging the gap between machine-learning output and a dynamic risk assessment more suitable for informing clinical decision-making. Using Bayesian reasoning (Payne et al., 2021), we demonstrate how to calibrate the level of informativeness of each data point. Specifically, we leveraged the odds formulation of Bayes’ Theorem, which offers a computationally elegant method for sequential updating. This method tracks seizure risk using the log-odds of a seizure event. The current log-odds are incrementally adjusted with each new data point using predictions derived from each one-second stride of the five-second ECoG data window. This updating mechanism involves a simple additive process in which the current log-odds are changed by adding the *log of the positive likelihood ratio* (LR+), which quantifies the diagnostic utility of the new prediction. The LR+ value is computed by applying the SVM predictor on held-out test data and computing the ratio of true preictal to true interictal for each bin of confidence predictions.

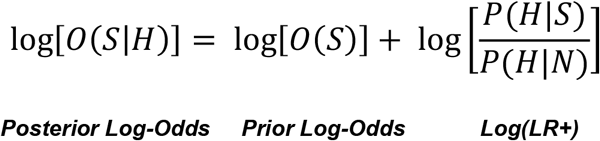

***Equation 1:*** *Bayesian updating method using the log-odds formulation. Odds are computed as the ratio of the probability of an event divided by the probability of that event not occurring, O = p/(1-p), which means that p = O/(O+1). Since the events are mutually exclusive and exhaustive, the denominator in the Bayes formula cancels out. The posterior log-odds can be computed as the prior log-odds plus the log of the positive likelihood ratio (LR+). The LR+ is the ratio of observing a prediction of that confidence level, given it really is a seizure, divided by the probability of such an observation if it were not. As each new prediction comes in, the running log-odds can be updated by adding the new log(LR+)*.

Bayesian updating has been found to be comparable to, and in some cases, superior to the firing power method, which also monitors the density of predicted samples within a given time window. The logarithms of the odds ratio on unseen test seizure data binned by confidence level were fitted with a linear function. This reflects an empirical measure of how determinative a point prediction should be considered towards the overall forecast.

### Radius of Gyration in Feature Space

The radius of gyration (R_g_) metric was used to obtain more rigorous quantification of the typical hypervolume explored in phase space. Defined as the root mean squared distance to the center of mass, the value for a collection of points in feature space is calculated as 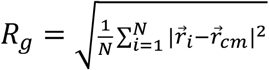 where the center of mass is 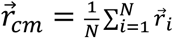 The R_g_ value is commonly employed to measure the compactness of folded proteins (Tanner, 2016). When applied to the interictal period, a larger R_g_ captures both a general expansion of the hypervolume explored, as well as the influence of rare, highly connected events, which significantly increase the radius of gyration for that time period.

## RESULTS

In figure 4, a Seaborn Pairplot illustrating network features at various thresholds for a single patient reveals distinct spatial organization. The core preictal period, defined as 1-4 minutes before seizure onset (red points), occupied a notably smaller, more constrained region of feature space compared to both intermediate preictal and interictal data points. Furthermore, after applying SMOTE and PCA preprocessing, a plot of the first and second principal components also demonstrated this spatial confinement of preictal points. The marginal histograms further highlight the key difference that the scale-invariant power-law behavior characteristic of healthy brain function, represented by a long tail, is present in the interictal data (blue) but is suppressed in the preictal data (green), instead having a more exponential cutoff that occurs when the system is far from the critical point (L. R. Nemzer et al., 2020).

**Figure 4.**
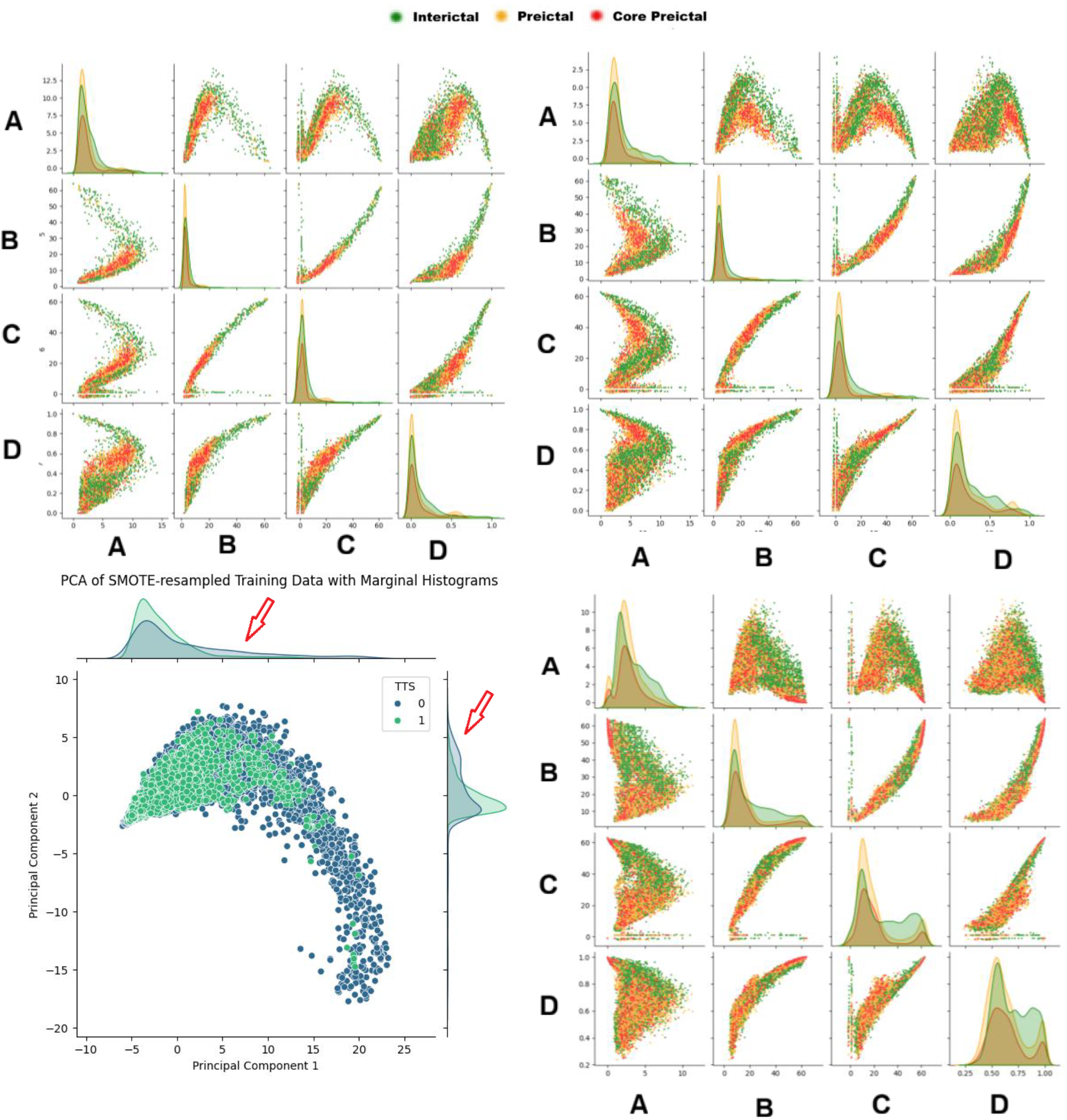
Pairplots of raw feature data at increasing thresholds showing the constriction of phase space explored during the core preictal and intermediate preictal periods compared with the interictal data. The rows and columns represent the four network features: (A) Normalized Node Degree Variance, (B) Size of Largest Clique, (C) First Eigenvalue, and (D) Average Clustering Coefficient. The distance thresholds are increasing clockwise from upper left, [0.250, 0.325, 0.500]. The data points were classified into core preictal (Red, 1-4 minutes prior to seizure onset), intermediate preictal (Orange, 4-10 minutes prior to seizure onset, masked in training data), and interictal (Green 10-59 minutes prior to seizure onset). Lower Left: Comparison of SMOTE-resampled PC1 and PC2 showing constriction of ictal (light green) compared with interictal (blue). Notice the heavy tail in the marginal histograms present in the interictal data that is suppressed in the preictal data (red arrows).

The characteristic suppression of phase space excursions is a key observation during the core preictal period, a phenomenon which is visually represented in Figure 5 (an animated version of which is provided in the **supplementary material** for dynamic viewing). In this figure, the x-axis represents the size of the largest clique, and the y-axis represents the average clustering coefficient, both measured at a threshold of 0.450. The red traces correspond to data sampled exclusively during the preictal phase, while the green traces represent data from the interictal period. Crucially, these red and green traces represent equal amounts of time elapsed, ensuring a fair temporal comparison of the system’s dynamic behavior in the two states. The background of the figure, rendered in blue, serves as a visual representation of the total extent of the feature space explored across the entire data block. The data from the preictal period is markedly sequestered compared with the interictal period. That is, the system’s trajectory during the preictal phase remains confined to a significantly smaller region of phase space and does not exhibit the extensive, wide-ranging excursions that are routinely observed and characteristic of the system’s behavior during the interictal period. This constriction in the dynamic movement within the feature space suggests a fundamental change in the underlying neuronal dynamics, shifting from a state of complex, high-dimensional exploration to one of constrained, low-dimensional organization that is more fragile to phase transitions as the system approaches seizure onset.

**Figure 5.**
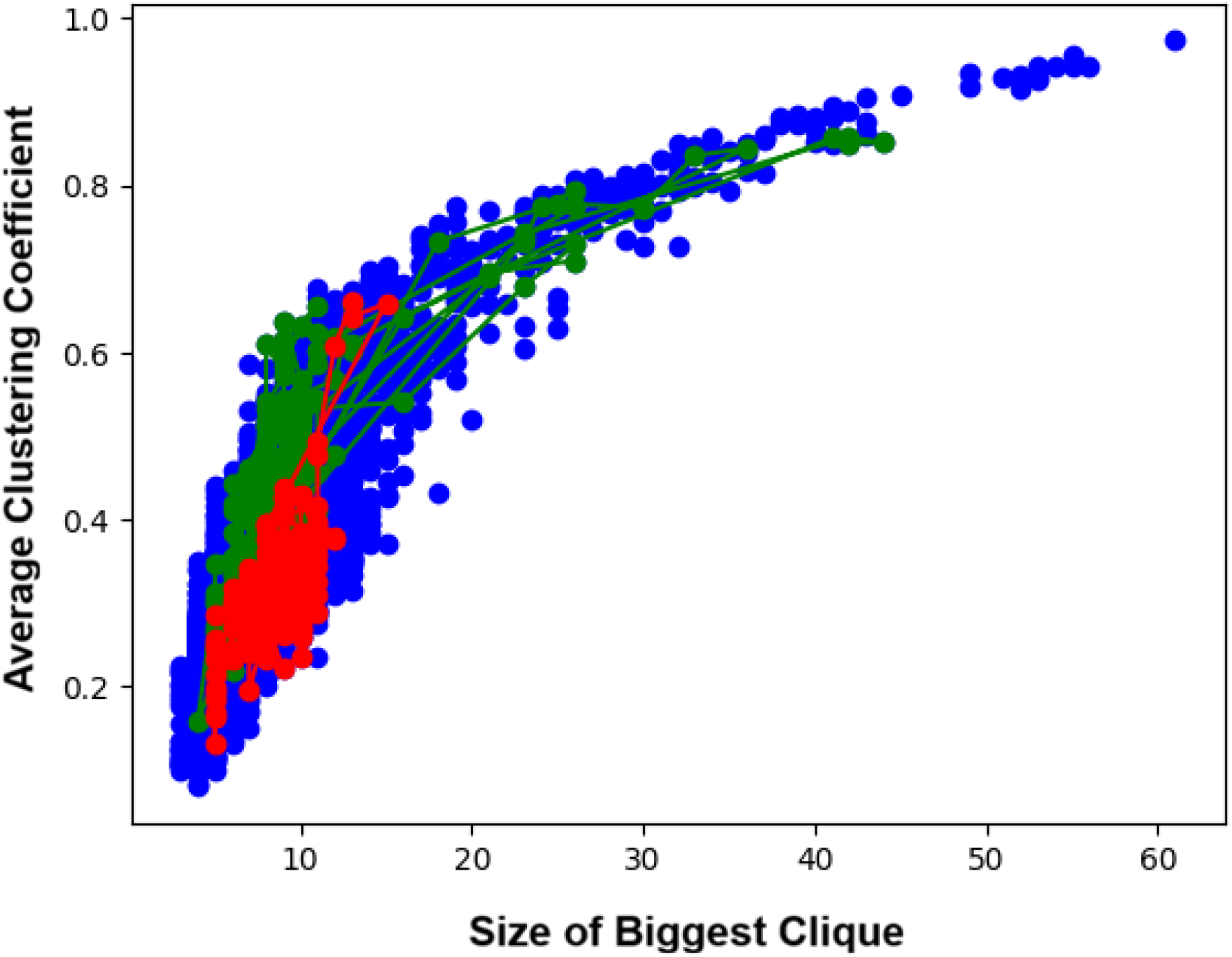
Phase space constriction shown by plotting the size of biggest clique on the x-axis and the average clustering coefficient on the y-axis at a threshold of 0.45. The blue data points represent the total feature space explored during the data block. The red (preictal) and green (interictal) traces represent equivalent elapsed time.

To quantify changes in the explored phase space, we standardized the size of the biggest clique and the average clustering coefficient for each of five patients to have a zero mean and unit variance. We calculated the mean radius of gyration for the collection of points identified as either inside or outside the core preictal time. Theoretically, the long-term average of the radius of gyration is expected to converge to the square root of the number of variables, which is approximately 1.41 in this case, since N = 2. The difference between the core preictal and other time periods was most pronounced at smaller distance thresholds for edges. As illustrated in Figure 6, this disparity diminished when the threshold for edge connection was increased.

**Figure 6.**
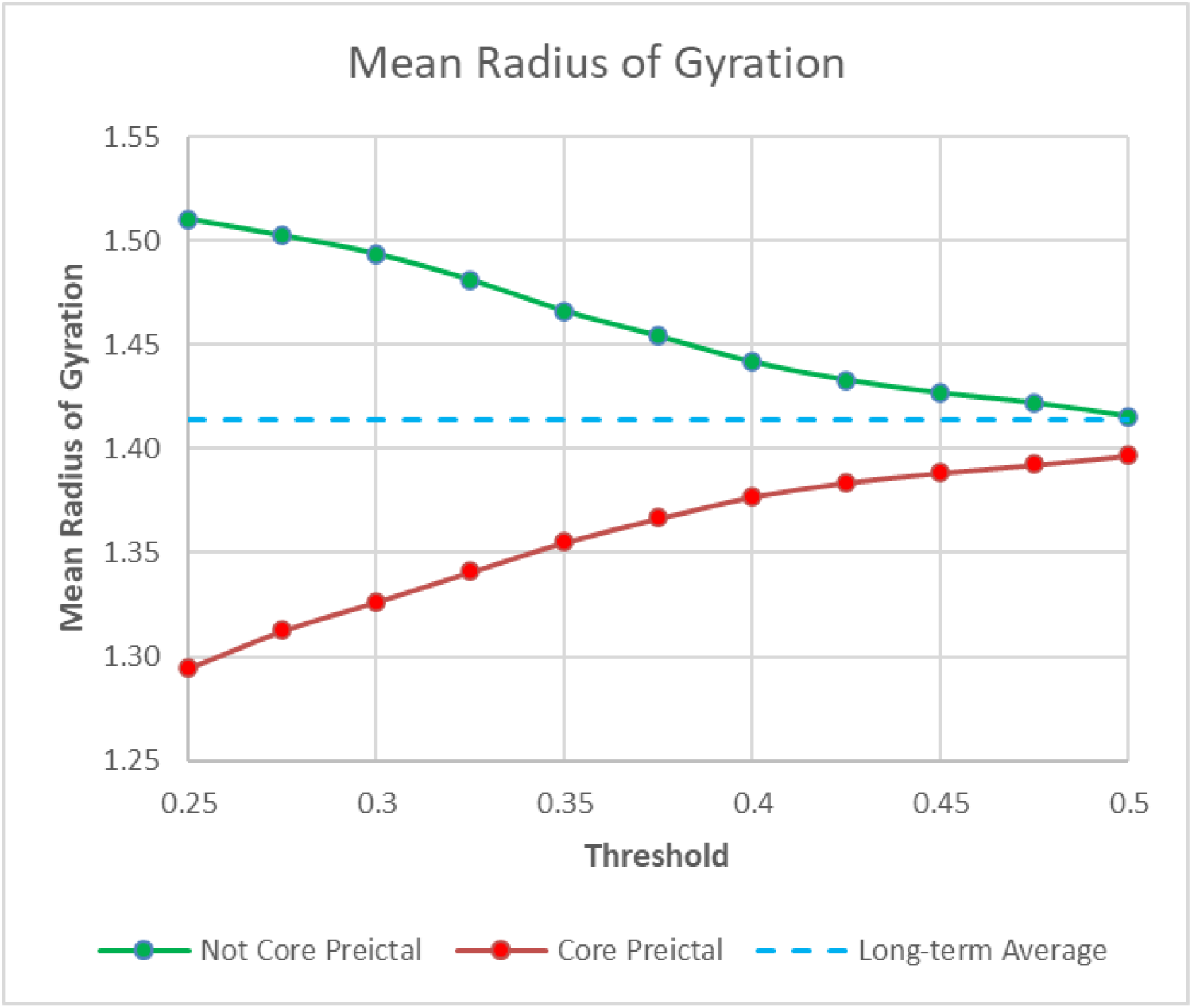
The mean radius of gyration by threshold across the parameter space defined by the size of the biggest clique and the average clustering coefficient for five patients. Notably, the core preictal phase (1-4 minutes, Red) demonstrates more compact exploration compared to other time periods (Green). The blue line illustrates the theoretical long-term average of approximately 1.41, which is the square root of the number of variables (N=2) that have been standardized to possess zero mean and unit variance.

The machine-learning predictions for individual datapoints were obtained with SVM. An example receiver operating curve (ROC) and confusion matrix are shown in figure 7. The training data was masked to remove the intermediate preictal data points, and only used core preictal (class 1) and interictal (class 0).

**Figure 7.**
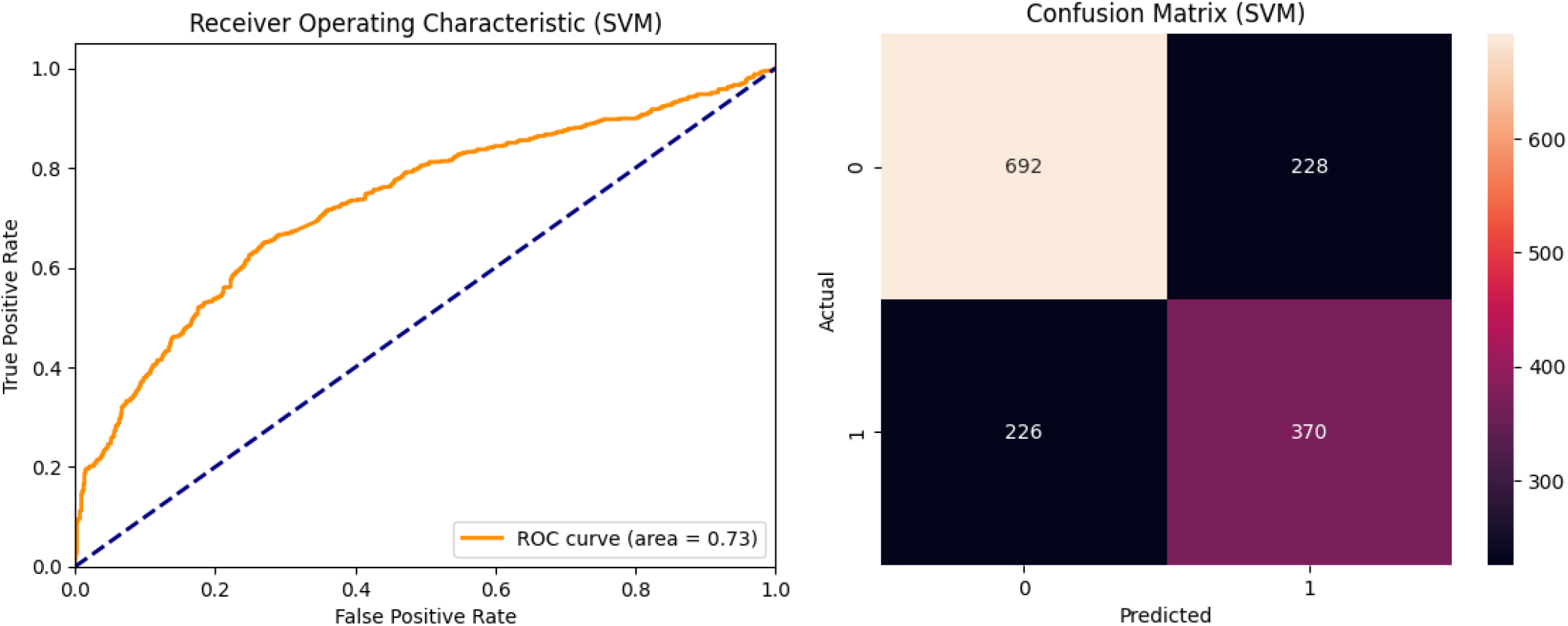
Receiver operator curve and confusion matrix for SVM. Accuracy: 0.7005. Sensitivity: 0.6208. F1 Score: 0.6198. ROC AUC: 0.7281

After the SVM predictor was trained, it was applied to held-out data of unseen seizure events from the test set, including intermediate interictal data. To calibrate the LR+, the predictions were binned by confidence (figure 8, top). The common logarithm of the ratio of true class of 1 versus true class of 0 was plotted as a function of confidence (figure 7, bottom). A linear fit was applied, with the constraint that a prediction 0.5 was considered to be uninformative, so LR+ = 1 and thus log(LR+) = 0. This constraint anchored the calibration curve, ensuring that the final LR+ values were properly scaled and centered around the point of maximal uncertainty. The resulting linear equation then provided a continuous mapping from any raw SVM confidence score to a calibrated, evidence-based value to update the overall Bayesian real-time risk forecast. Due to the small number of observations at extreme confidence levels, only values between 0.3 and 0.9 were used, and other predictions were set to zero.

**Figure 8.**
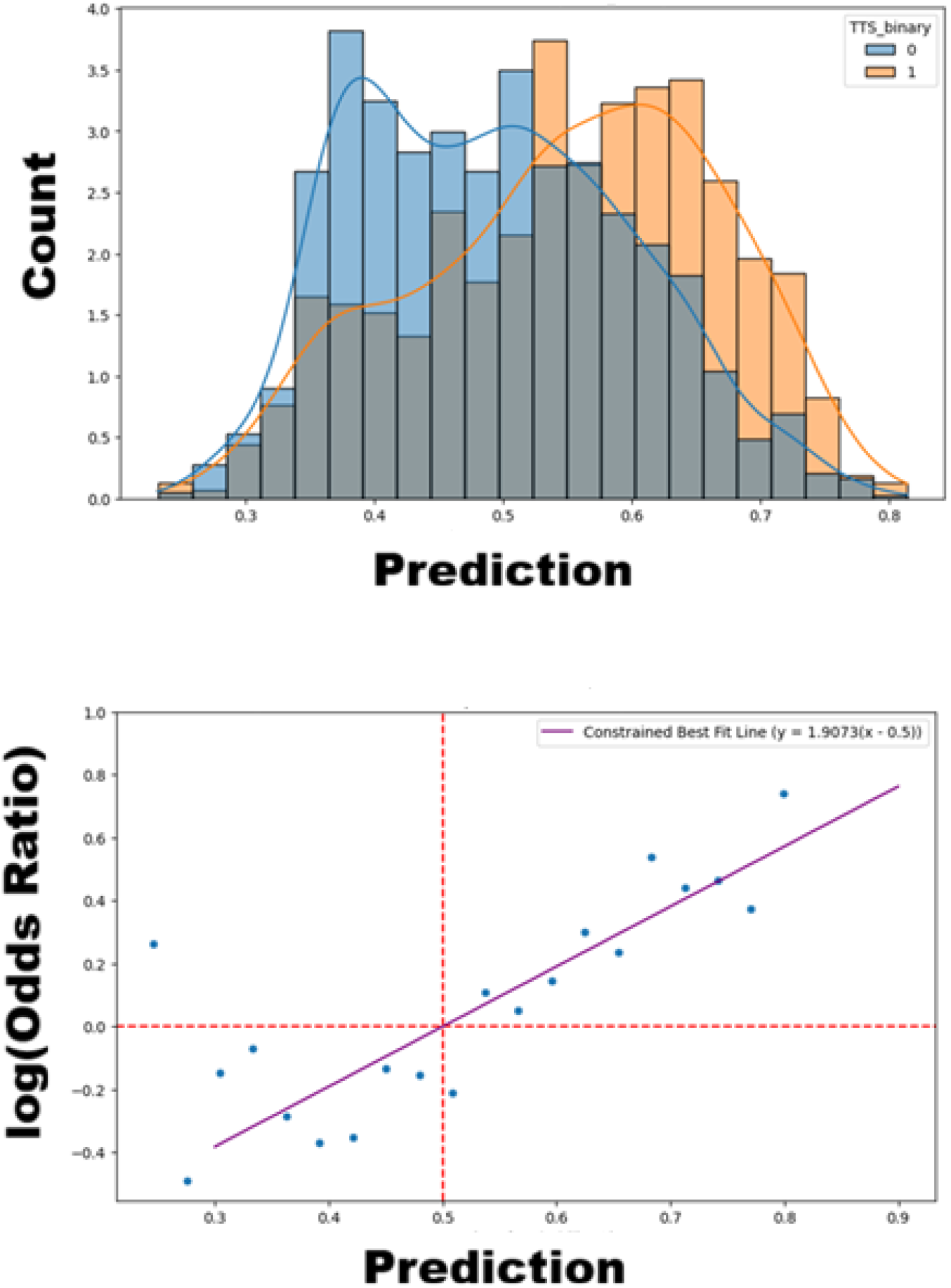
Calibrating log odds. Binned predictions of test data (top). Linear fit passing thorough (0.5,0) representing the informative value of each observation (bottom). Due to the small number of observations in the extremes, the range used was from 0.3 to 0.9 and zero updates outside that.

As a proof of concept, the algorithm was applied to stitched data combining different data blocks. Figure 9 (top) shows the time to seizure, and figure 8 bottom shows the corresponding real-time seizure risk forecast.

**Figure 9.**
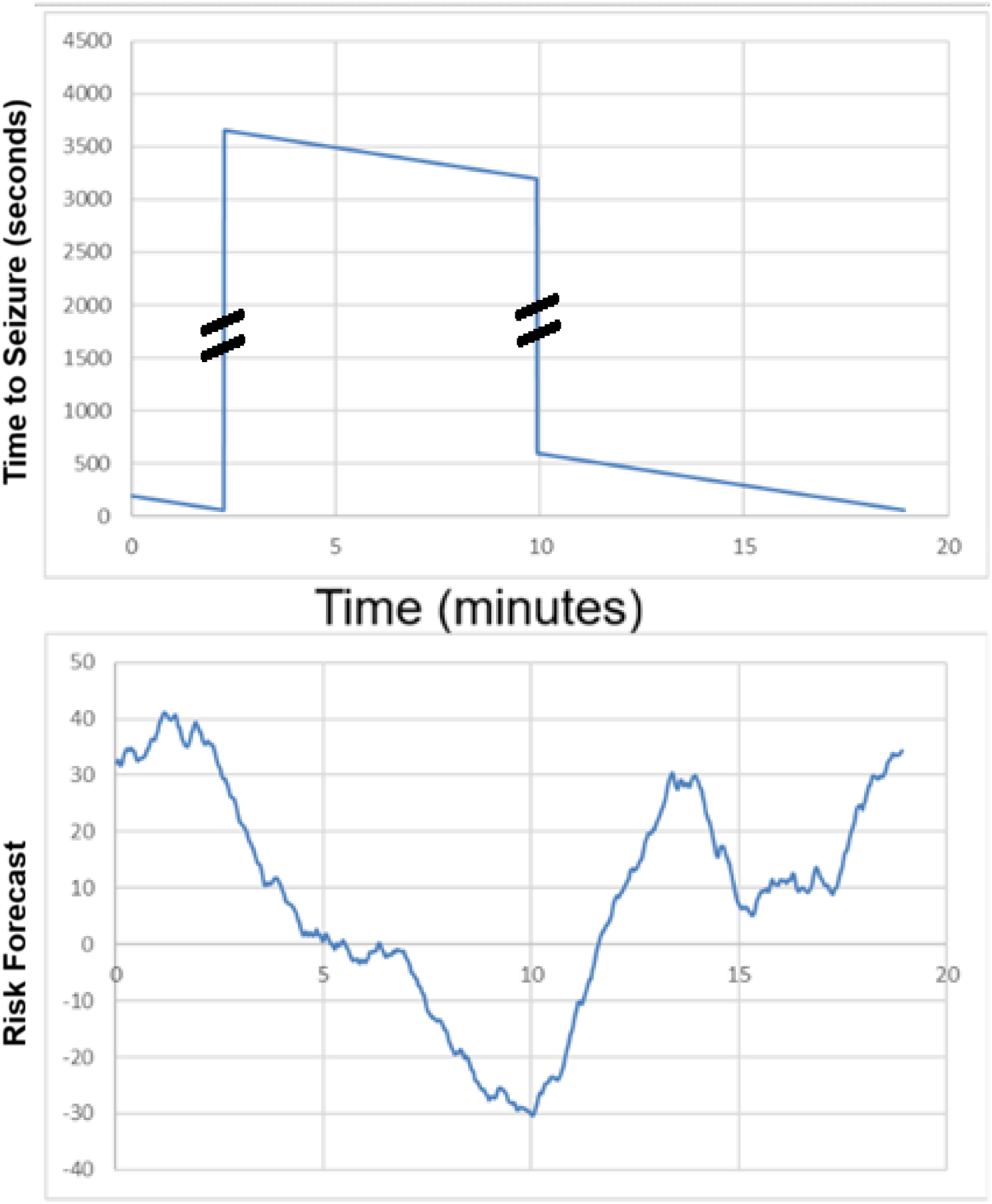
Proof of concept real-time seizure risk forecast using stitched data from different data blocks. Time to seizure in seconds (top). Corresponding real-time Bayesian real-time risk forecast (bottom).

## DISCUSSION

While the results of this study do not rule out the possibility of longer “preictal periods” based on changes in biomarkers or connectivity patterns, which could offer indicators significantly longer in advance, they strongly imply that the most reliable, short-term prediction window is much smaller than what has been traditionally assumed. Therefore, the effective working definition of when a seizure is predictably imminent, based on electrophysiological changes, may require revision to a narrower time frame. Further research is necessary to fully characterize the neurophysiological signature of the approximately four-minute “fragile” state and explore its critical implications for developing highly accurate, real-time seizure warning systems. It is important to emphasize that the findings here may vary significantly across patients, or even among seizure events for the same patient. More investigation is needed to establish the generalizability of these results.

## CONCLUSION

We propose a computationally efficient methodology for forecasting real-time seizure risk using intracranial electrocorticographic data. This approach is highly adaptable to individual needs, allowing patients to personalize the alert threshold. This personalization enables a balance between maximizing sensitivity (reducing missed seizures) and minimizing the false-positive rate (decreasing unnecessary alerts). The findings of this work could help point the way for the development of wearable devices capable of alerting both patients and healthcare providers, with the methodology suitable for implementation on small electronic devices or smartphone applications. Our approach represents progress towards lightweight, interpretable physiological forecasting for seizure prediction, informed by the critical brain hypothesis. A crucial finding that enhances the clinical relevance of our model is the absence of large excursions within feature space representing long-distance synchronization immediately before seizure onset. This lack of long tail deviations during the critical preictal transition, which are present during healthy brain function, could be physiologically relevant. This outcome aligns with the hypothesis that the transition to a seizure state is preceded by a subtle reorganization of brain network activity that loses resilience against a transition to pathological synchronization, rather than an immediate breakdown. Therefore, the interpretability of our model is rooted in tracking these changes, which offers a more stable and physiologically plausible foundation for reliable risk forecasting.

## Supporting information

Figure 4 Animation

Figure 5 Animation

## ACKNOWLEDGMENT

This work was supported by NSU PRG #334908. The authors would like to express their appreciation to Dr. Francis Motta for his helpful discussions.

## CONFLICT OF INTEREST STATEMENT

The authors report no conflicts of interest.

## Notes

https://campus-technologies.de/en/the-european-epilepsy-database-2/

### Competing Interest Statement

The authors have declared no competing interest.

