## Supplementary figures and images for "Electrocorticographic Network Feature Space Constriction as a Preictal Biomarker"

### Figure 4 Animation

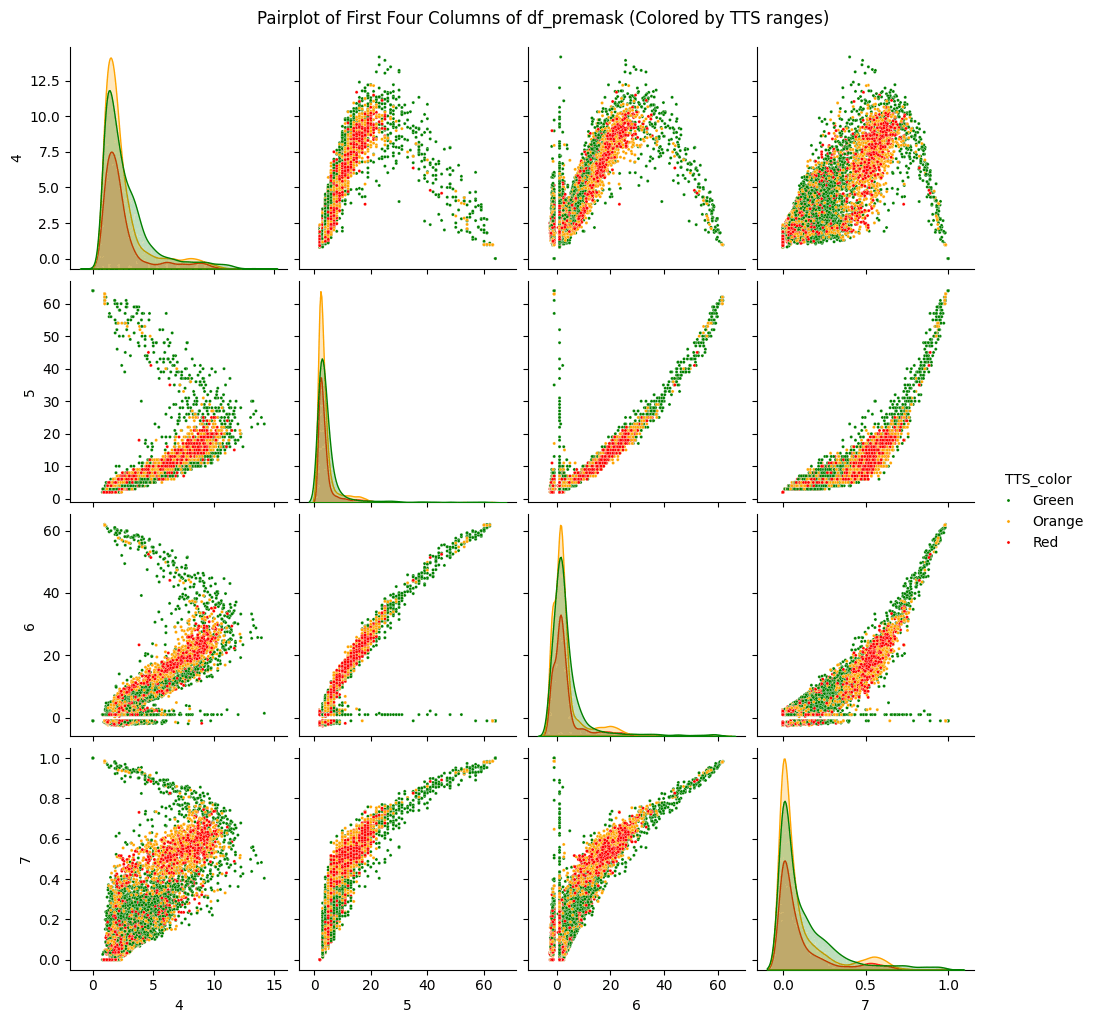

### Figure 5 Animation

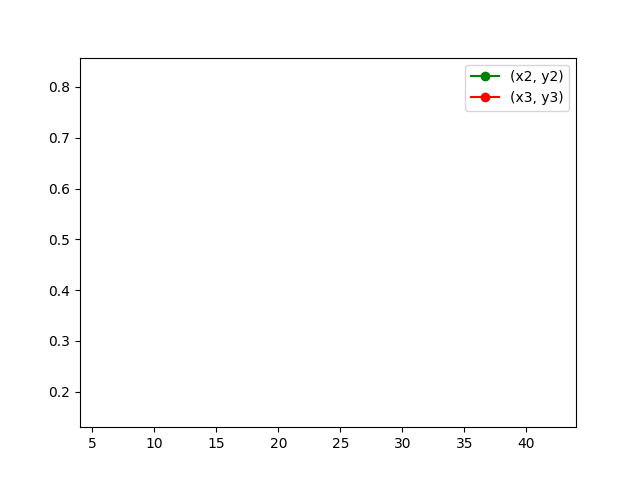
